# Usutu virus escapes langerin-induced restriction to productively infect human Langerhans cells, unlike West Nile virus

**DOI:** 10.1101/2021.08.17.456611

**Authors:** Marie-France Martin, Ghizlane Maarifi, Hervé Abiven, Marine Seffals, Nicolas Mouchet, Cécile Beck, Charles Bodet, Nicolas Lévèque, Nathalie J. Arhel, Fabien P. Blanchet, Yannick Simonin, Sébastien Nisole

## Abstract

Usutu virus (USUV) and West Nile virus (WNV) are phylogenetically close emerging arboviruses transmitted by mosquitoes, and constitute a global public health threat. Since USUV and WNV enter the body through the skin, the first immune cells they encounter are skin-resident dendritic cells, the most peripheral outpost of immune defense. This unique network is composed of Langerhans cells (LCs) and dermal DCs, which reside in the epidermis and the dermis, respectively.

Using human skin explants, we show that while both viruses can replicate in keratinocytes, they can also infect resident DCs with distinct tropism, since WNV preferentially infects DCs in the dermis, whereas USUV has a greater propensity to infect LCs. Using both purified human epidermal LCs (eLCs) and monocyte derived LCs (MoLCs), we confirm that LCs sustain a faster and more efficient replication of USUV compared with WNV and that this correlates with a more intense innate immune response to USUV compared with WNV. Next, we show that ectopic expression of the LC-specific C-type lectin receptor (CLR), langerin, in HEK293T cells allows WNV and USUV to bind and enter, but supports the subsequent replication of USUV only. Conversely, blocking or silencing langerin in MoLCs or eLCs made them resistant to USUV infection, thus demonstrating that USUV uses langerin to enter and replicate in LCs. Altogether, our results demonstrate that LCs constitute privileged target cells for USUV in human skin, because langerin favors its entry and replication. Intriguingly, this suggests that USUV efficiently escapes the antiviral functions of langerin, which normally safeguards LCs from most viral infections.

## Introduction

Usutu virus (USUV) and West Nile Virus (WNV) are emerging arboviruses belonging to the *Flavivirus* genus of the *Flaviviridae* family, which comprise many other human pathogenic viruses, including Zika virus (ZIKV), Dengue virus (DENV) or Yellow fever virus (YFV). They are phylogenetically close and both belong to the Japanese encephalitis virus (JEV) antigenic complex ^1^. WNV and USUV have recently expanded outside of Africa, where they both originate: WNV is now endemic throughout much of the world whereas the spread of USUV has recently dramatically increased in Europe and it has become endemic in several European countries ^2–6^. WNV has caused several outbreaks among humans and birds, whereas USUV has been mainly involved in major avian epizootics in Europe and has been associated with neurological disorders in humans ^3–5,7,8^. Therefore, these two viruses are considered as serious potential threats to human and animal health.

USUV and WNV are maintained in the environment through an enzootic cycle involving mosquitoes (mainly of the genus *Culex*) and birds ^9^. Infected mosquitoes can incidentally transmit the viruses to mammals, such as human and horses, which are dead-end hosts but can both develop severe neurological disorders. Since viruses are directly inoculated by the bite of an infected mosquito, the first organ to get infected is the skin.

The skin is a complex organ, composed of two main layers: the dermis, made up of connective tissue produced by fibroblasts, and the epidermis, a multilayered stratified epithelium mostly constituted of keratinocytes. The immune surveillance of this most peripheral organ of the body is mainly ensured by a unique network of dendritic cells (DCs), composed of Langerhans cells (LCs), which is the only DC subset that resides in the epidermis, and dermal DCs (dDCs), which are mostly located in the upper dermis ^10,11^. LCs and dDCs are specialized in the recognition and capture of pathogens in the skin. Following antigen uptake and processing, they migrate to local draining lymph nodes in order to activate effector T cells. The capacity of skin-resident DCs to capture various pathogens is conferred by the expression of specific pattern recognition receptors (PRRs), including C-type lectins receptors (CLRs), which bind carbohydrate structures associated to viruses, fungi or bacteria ^12^. Among the CLRs, langerin (or CD207) is exclusively expressed by LCs in humans ^13^. Langerin is not only present at the cell surface but also within rod-shaped cytoplasmic organelles with a striated appearance, termed Birbeck granules (BGs) ^13^. These are subdomains of the endosomal recycling compartment that are involved in pathogen degradation and antigen processing ^13,14^. The role of langerin in the capture and degradation of viruses by LCs has been particularly well described in the case of HIV-1 ^15,16^. Another important CLR, mostly expressed by DCs, including dDCs, is DC-SIGN (or CD209) ^17^. Although CLRs have clear antiviral functions, many viruses are able to hijack these receptors to their advantage. In particular, DC-SIGN has been shown to bind HIV-1 gp120 and promote efficient *trans*-infection of T cells ^18^. DC-SIGN can also be used by many viruses to infect immature DCs, such as Cytomegalovirus (CMV) ^19^ or Ebola virus (EBOV) ^20^ but also several flaviviruses, including DENV and WNV ^21–23^. The viral hijacking of langerin seems to be much rarer and has only been formally demonstrated for influenza A virus (IAV), although it was not determined whether LCs constitute target cells ^24^. In contrast, some viruses have been shown to infect Langerhans cells in human skin, including DENV ^25–27^.

The role of skin cells in WNV infection, amplification and spread has been studied both *in vitro* and *in vivo* ^6,28,29^. In particular, it has been shown that infected mosquitoes inject up to 10^6.6^ PFU WNV particles into the skin, mostly outside blood vessels ^30^, thus promoting the infection of skin cells, including keratinocytes ^31^. WNV was shown to infect monocyte-derived DCs ^22,32–34^, but it not known whether it can infect skin-resident DCs. Although LCs were shown to migrate to local lymph nodes following cutaneous infection with WNV, it was not determined whether these were infected or not ^35^. Similarly, since USUV has received less attention than WNV, its cellular tropism in the skin has not yet been investigated and its ability to infect skin-resident DCs is entirely unknown.

In this study, we evaluated the capacity of USUV and WNV to infect human skin-resident DCs, including LCs. Using both human primary skin-isolated and monocyte-derived LCs (MoLCs), we report that, although WNV is taken up by LCs to some degree, USUV enters and replicates within LCs much more efficiently than WNV. In particular, we show that human LCs support productive infection of USUV and constitute privileged target cells for this virus. The innate immune response triggered by USUV was also much more intense than that by WNV. Finally, we show that while both USUV and WNV can enter cells following their interaction with langerin, only USUV escapes langerin-induced restriction in order to replicate in Langerhans cells.

## Material and Methods

### Cells

C6/36 cells (CRL-1660), Vero (CCL-81) and HEK293T (CRL-11268) were purchased from the American Type Culture Collection (ATCC). STING-37 reporter cells were kindly provided by Pierre-Olivier Vidalain (CIRI, Lyon, France). All cells were cultured in Dulbecco’s modified Eagle’s medium supplemented with 10% fetal bovine serum (Serana), 1% Penicillin/Streptomycin (Gibco) and maintained in 5% CO2 at 37°C, except C6/36 cells which were grown at 28°C.

### Viruses

USUV Europe 2 (TE20421/Italy/2017) was kindly provided by Giovanni Savini (Istituto Zooprofilattico Sperimentale dell’ Abruzzo e del Molise “G. Caporale”, Teramo, Italy). USUV Africa 2 (Rhône 2705/France/2015) and WNV lineage 2 (WNV-6125/France/2018) were provided by ANSES (National Agency for Food, Environmental and Occupational Health Safety, France). The lineage 1 clinical strain of WNV was isolated from a human brain during the epidemic that occurred in Tunisia in 1997 and was provided by Isabelle Leparc-Goffart (French National Reference Center on Arboviruses, Marseille, France). The origin and history of viral isolates used in the study are summarized in Supp. Table 1. Viral strains were amplified on C6/36 cells. Supernatants were collected 5 days after infection and their titers were determined on Vero cells, using the Spearman-Karber method, and expressed as TCID50/ml ^36^. All infections were performed by incubating cells with the virus in serum-free medium for 2 h, before the cells were washed and incubated with medium supplemented with 10% FBS. In the case of short infections (30 min), the cells were washed at the end of the incubation time.

### Antibodies and reagents

The primary antibodies used were mouse anti-dsRNA (J2 clone, Scicons), mouse pan-flavivirus (4G2 clone, Novus Biologicals), rabbit anti-MX1 (Thermo Fischer Scientific), mouse APC conjugated anti-langerin (clone 10E2, BioLegend), mouse anti-langerin (clone D9H7R, Cell Signaling), recombinant anti-DC-SIGN (clone REA617, Miltenyi Biotec), mouse anti-CD1a (Novus Biologicals), FITC-conjugated anti-CD1a (clone HI149, Miltenyi Biotec), FITC-conjugated anti-HLA-DR (clone AC122, Miltenyi Biotec), mouse anti-HSP90 (clone F-8, Santa Cruz), and mouse anti-GAPDH (clone 6C5, Merck Millipore). Secondary antibodies were goat anti-mouse AF488 (Thermo Fisher Scientific), donkey anti-mouse AF568 (Thermo Fisher Scientific), donkey anti-rabbit AF647 (Thermo Fisher Scientific) and goat anti-mouse or anti-rabbit HRP conjugates (GE Healthcare).

Mannan from *Saccharomyces cerevisiae* was purchased from Sigma-Aldrich.

### Plasmids and transfections

All ectopic transfections were performed using FuGENE® 6 Transfection Reagent (Promega) according to the manufacturer’s instructions. Plasmids expressing human langerin and DC-SIGN have been described previously ^15,37^. Expression of langerin and DC-SIGN constructs was assessed by flow cytometry, western blot, or indirect immunofluorescence. For siRNA transfection, non-targeting control siRNA (siRNA CTR) and siRNA specific for langerin (siRNA langerin) were purchased from Dharmacon as SMARTpools. Transfections were performed using DharmaFECT 4 (Dharmacon), as previously described ^38^.

### Blood samples, isolation and culture of human primary cells

Buffy coats from healthy donors were obtained from the Etablissement Français du Sang (EFS, Montpellier, France). PBMCs were isolated by density centrifugation using Lymphoprep medium (STEMCELL Technologies). CD14+ monocytes were isolated from PBMCs using CD14 MicroBeads (Miltenyi Biotec) and subsequently differentiated into MoDCs or MoLCs. Briefly, MoDCs were generated by incubating purified monocytes in Iscove’s Modified Dulbecco’s Medium (IMDM) supplemented with 10% FBS, 1% P/S, 2 mM L-glutamine, 10 mM Hepes, 1% non-essential amino-acids, 1mM sodium pyruvate and cytokines GM-CSF (Granulocyte-Macrophage Colony Stimulating Factor, 500 IU/ml) and IL-4 (500 IU/ml), both from Miltenyi Biotec (Cytobox Mo-DC). MoLCs were generated as MoDCs, except for the addition of 10 ng/ml of TGF-β (Peprotech) within the differentiation medium. Immature MoDCs and MoLCs were harvested at day 6 and cell differentiation was estimated by measuring the expression of DC-SIGN, langerin, CD1a and HLA-DR (class II) by flow cytometry (Supp. Figure 1A).

### Isolation of epidermal LCs from human skin explants

After the provision of fully informed consent, skin samples were obtained from patients undergoing abdominoplasty or mammoplasty plastic surgery at the Poitiers University Hospital, France. The use of all human skin samples for research studies was approved by the Ethics Committee (committee for the protection of persons) Ouest III (project identification code: DC-2014-2109).

Human skin samples processing was adapted from the method previously described in ^38^. Briefly, skin sheets were cut into 1 cm^2^ pieces and incubated with agitation in shaking water bath (at 175 strokes/minute) in RPMI medium (Gibco) containing collagenase A (1 mg/ml, Roche), DNase I (20 U/ml, Sigma, D4263-5VL) and Dispase II (1 mg/ml, Roche) overnight at 37°C, after which the epidermis was mechanically separated from the dermis using forceps. Epidermal sheets were cultured separately in RPMI with 10% human AB serum (Thermo Fisher Scientific) and 1% Penicillin/Streptomycin/Amphotericin B solution for 48 h, after which migratory cells were collected from media. Migratory cells were either used directly for experiments or subjected to a positive selection using CD1a Microbeads (Miltenyi Biotec), in order to enrich epidermal Langerhans cells. The expression of CD207 and HLA-DR (class II) was assessed by flow cytometry. Staining was performed on both total epidermal cells and following purification of CD1a+ cells, in order to estimate the enrichment rate (Supp. Figure 1B).

### Infection of human skin explants and multiplex immunofluorescence assay

The epidermal layer of 1 cm^2^ intact human skin explants (epidermis + dermis) was gently pricked using a fine hypodermic needle (Terumo Agani needle 25G, 0.5×25 mm) and the viral inoculum was applied on the pierced epidermis, in order to allow viruses to spread inside the tissue. USUV and WNV infections were systematically performed by the same experimenter and on explants from the same donor, following a standardized procedure, to ensure equal infection between explants. The pieces of skin were infected with 10^7^ TCID50/ml of USUV AF2 or WNV L1 in 500 µL of RPMI containing 2% human AB serum and 1% of P/S to submerge the tissue. After 4 h at 37°C, the viral inoculum was washed and the skin explants were incubated with medium supplemented with 10% human AB serum at 37°C. At 24 h post-infection, skin pieces were fixed in 4% paraformaldehyde (PFA) for 24 h at 4°C, and dehydrated in 70° ethanol. Samples were sent to the H2P2 platform (University of Rennes, France) to perform multiplex immunofluorescence assays. Paraffin-embedded tissue was cut at 4 µm, mounted on Adhesion slides (TOMO) and dried at 58°C for 12 h. Immunofluorescent staining was performed on the Discovery ultra-Automated IHC stainer, using the Discovery Rhodamine kit and the Discovery FAM kit (Ventana Medical Systems). Following deparaffination with Ventana Discovery Wash solution (Ventana Medical Systems) at 75 °C for 8 min, antigen retrieval was performed by using Tris-based buffer solution CC1 (Ventana Medical Systems) at 95°C to 100°C for 40 min. Endogen peroxidase was blocked with Disc inhibitor (Ventana Medical Systems) for 8 min at 37°C. After rinsing with reaction buffer (Ventana Medical Systems), slides were incubated at 37°C for 60 min with the pan-flavivirus antibody. After rinsing, signal enhancement was performed using anti-mouse HRP antibody (Ventana Medical Systems) incubated for 16 min and the discovery Rhodamine kit (Ventana Medical Systems). Following denaturation with Ventana solution CC2 (Ventana Medical Systems) at 100°C for 8 min, slides were incubated at 37°C for 60 min with the anti-CD1a antibody. After rinsing, signal enhancement was performed using anti-mouse HRP antibody (Ventana Medical Systems) incubated for 16 min and the discovery FAM kit (Ventana Medical Systems). Slides were then counterstained for 4 min with DAPI and rinsed. After removal from the instrument, slides were manually rinsed and placed on coverslips. Images were acquired on a confocal scanner. Quantifications were performed with the image analysis platform Halo (V3.2.1851.371), using the HighPlex module (V4.04). Whole sections were analyzed (representing approximately 100,000 cells per condition). The number of cells positive for CD1a and pan-flavivirus staining was determined in the epidermis and in the dermis.

### Viral titration (TCID50/mL)

According to the Centre National de Référence des Arboviroses protocol (IRBA, Marseille), Vero cells were plated in DMEM 2% FBS 1% P/S on a 96 well plate (in duplicate for each condition and for each independent experiment). Few hours later when cells were adherent and subconfluent, 100 µL of supernatant from previously infected cells per replicate was serially diluted in DMEM 2% FCS 1% P/S. Vero culture medium was removed and serial dilutions were applied on Vero plated wells (100 µL per well, 6 wells per serial dilution). On each plate a column was dedicated to control media (NI) and wells on the borders were filled with PBS to avoid evaporation. Infected plates were incubated 7 days at 37°C with 5% CO2. Cytopathic effects (CPE) were then evaluated using an optical microscope and titers were calculated in TCID50/mL using the Spearman-Karber method (Karber G. *et al*, 1931).

### Quantification of secreted cytokines and chemokines

Total IFN secreted by monocytes, MoDCs and MoLCs was titrated on STING-37 reporter cells, which correspond to HEK293 cells stably expressing an IFN-stimulated response element (ISRE)-luciferase reporter gene ^39^. A standard curve was established by applying known titers of recombinant IFN-α2a (R&D Systems) onto STING-37 cells. Luciferase induction in STING-37 cells was determined using the Bright-Glo reagent (Promega), according to manufacturer’s instructions, and luminescence signal was acquired on a TECAN Infinite 200. Quantification of cytokine and chemokine levels in culture media was performed using the LEGENDPlex kit from BioLegend (human anti-virus response panel), according to the manufacturer’s recommendations.

### Immunofluorescence assays

Cells were plated on poly-D-lysine coated coverslips, fixed with 4% PFA (Alfa Aesar) for 10 min, permeabilized with 0.1% Triton X100 for 15 min, neutralized with 50 mM NH4Cl for 10 min, and blocked with 2% BSA for 10 min. Cells were incubated with primary and secondary antibodies for 1 h and 45 min, respectively, at room temperature in a wet chamber. Finally, cells were labelled with Hoescht and mounted in SlowFade antifade reagent (Thermo Fischer Scientific). Images were acquired with a Leica SP5-SMD confocal microscope. Mander’s coefficients were determined by counting 3 fields of around 300 cells per condition using the JAcoP plugin (ImageJ).

### Flow cytometry analysis

All cells were fixed with 2% PFA for 30 min and permeabilized in a PBS/1% BSA/0.05% saponin solution for 30 min prior to intracellular staining with corresponding primary antibodies for 1h at 4°C diluted in the permeabilization solution and then incubated with the corresponding secondary antibody for 30 min. For flow cytometry analysis, all acquisitions were done with Fortessa cytometer (B Becton Dickinson D), data were collected with FACSDiva software (Becton Dickinson) and were processed with FlowJo software (Treestar Inc.).

### Immunoprecipitation

HEK293T cells were transfected with an empty plasmid (pcDNA3.1) or with a plasmid encoding langerin and infected 24 h post-transfection with USUV AF2 and WNV L1 at MOI 5. 30 minutes after infection, cells were lysed in IP lysis buffer (50 mM Tris-HCl (pH7.4), 150 mM NaCl, 1 mM EDTA, 1% Triton X-100, 20 mM N-ethylmaleimide, EDTA-free protease inhibitor cocktail (Roche)) for 1 h at 4°C. Cell lysates were then incubated overnight at 4°C with pan-flavivirus anti-E antibody and protein G Sepharose beads (Thermo Fisher Scientific). Beads were washed three times and eluted with 2× SDS loading buffer before Western blot analysis.

### Western Blot

For the detection of the viral envelope protein, the anti-pan-flavivirus antibody was used under non-reducing and non-denaturing conditions. Briefly, cells were lysed in buffer containing Tris pH 7.6 1mM, NaCl 150mM, Deoxycholate 0.1% (deoxycholic acid), EDTA 1mM and Triton 1% during 30 min at 4°C and then Laemmli 2X with SDS but without β-mercaptoethanol. Cell lysates were loaded on 10% ProSieve gel (LONZA, LON50618), then subjected to electrophoresis. Chemiluminescent acquisitions were done on a Chemidoc™ MP Imager and analyzed using Image Lab(tm) desktop software (Bio-Rad Laboratories).

### RT-qPCR

Total RNA was extracted using a RNeasy Mini kit (Qiagen) following the manufacturer’s instructions. RNA concentration and purity were evaluated by spectrophotometry (NanoDrop 2000c, Thermo Fischer Scientific). A maximum of 500 ng of RNA were reverse transcribed with both oligo dT and random primers using a PrimeScript RT Reagent Kit (Perfect Real Time, Takara Bio Inc.) in a 10 µL reaction. Real-time PCR reactions were performed in duplicate using Takyon ROX SYBR MasterMix blue dTTP (Eurogentec) on an Applied Biosystems QuantStudio 5 (Thermo Fischer Scientific). Transcripts were quantified using the following program: 3 min at 95°C followed by 40 cycles of 15 s at 95°C, 20 s at 60°C and 20 s at 72°C. Values for each transcript were normalized to expression levels of RPL13A (60S ribosomal protein L13a), using the 2-ΔΔCt method. Primers used for quantification of transcripts are indicated within Supp. Table 2.

## Results

### USUV has the propensity to infect human epidermal Langerhans cells (eLCs)

First of all, we compared the capacity of USUV and WNV to infect DCs within human skin, using a strain of USUV Africa 2 (USUV AF2) isolated in France in 2018, and a clinical strain of WNV belonging to the lineage 1 (WNV L1). Explants were infected with USUV AF2 or WNV L1, and paraffin-embedded tissues were analyzed by immunofluorescent staining using an anti-CD1a antibody and a pan-flavivirus antibody targeting the envelope protein E of flaviviruses, to label skin-resident DCs and infected cells, respectively (Figure 1A). As for WNV, most USUV-infected cells were epidermal CD1a-negative, thus suggesting that keratinocytes are also the main targets of USUV. However, a substantial proportion of double-positive cells could also be observed, mostly in the case of USUV infection (Figure 1A). We performed a quantification of CD1a-positive infected cells both in the epidermis and dermis, which confirmed that the proportion of double-positive cells was approximately 2-times higher with USUV compared to WNV (Figure 1B). Interestingly, CD1a-positive cells infected by USUV were almost exclusively found in the epidermis, suggesting that USUV preferentially infects LCs. In contrast, WNV was found to infect indifferently epidermal (eLCs) and dermal (dDCs) DCs (Figure 1B). In order to confirm these observations, we infected total epidermal cells or purified eLCs from human skin explants with USUV or WNV for 24 h at MOI 2 (Supp. Figures 1B and 1C). As with intact human skin, USUV was found to infect epidermal cells more efficiently than WNV and strikingly, was able to infect nearly all eLCs (Figure 1C). Furthermore, since we labeled the cells with an antibody targeting dsRNA, which recognizes replicating viral genomes, our results suggest that both viruses can replicate within LCs, but that USUV infection is more efficient (Figure 1C). This was confirmed by immunofluorescence staining of purified human epidermal cells, which allowed us to detect USUV replicating within LCs (Figure 1D). Some replicating WNV could be detected, but at very low levels (Figure 1D). Similarly, viral RNA was quantified by RT-qPCR and further confirmed that the amount of USUV in purified epidermal LCs was approximately 10 times higher than that of WNV (Figure 1E). Altogether, our results show that in human skin, both WNV and USUV can infect epidermal CD1a-cells (mainly keratinocytes), as well as eLCs and dDCs. USUV however, was more efficient than WNV to infect skin-resident DCs and in particular eLCs.

**Figure 1.**
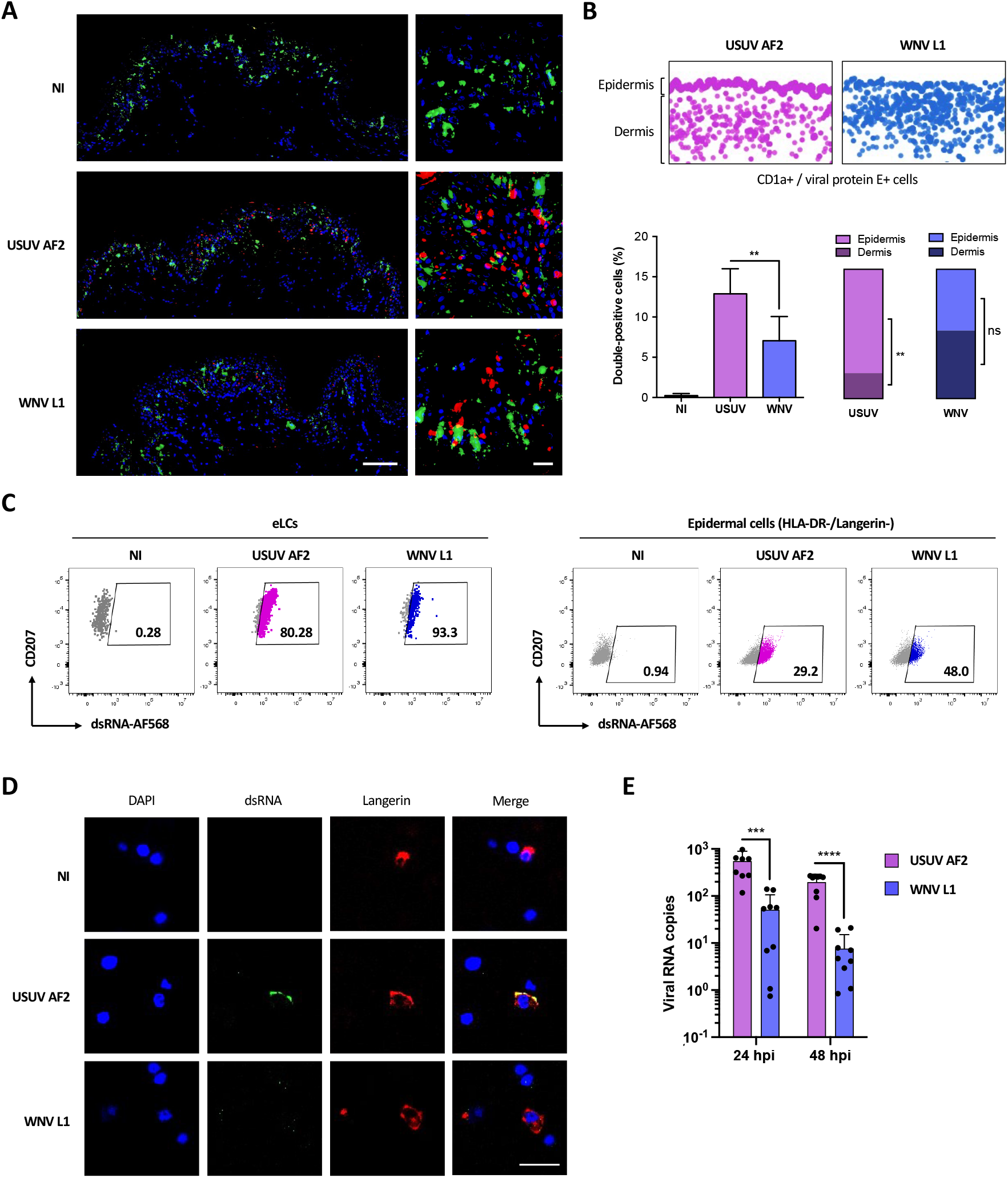
Unlike WNV, USUV infects preferentially Langerhans cells in human skin. **(A)** Human skin explants were left uninfected (NI) or infected with 10^7^ TCID50/ml of USUV AF2 or WNV L1. 24 h post-infection, tissues were fixed and paraffin-embedded, and analyzed by multiplex immunofluorescent assay using a pan-flavivirus anti-viral envelope E antibody (red) and an anti-CD1a antibody (green). Nuclei were stained with DAPI (blue). Images were acquired on a confocal scanner. Scale bars: 100 µm (left) or 20 µm (right). **(B)** Pan-flavivirus (red) and CD1a (green) staining from (**A**) was quantified using the image analysis platform Halo. Screenshots of double positive cells (infected CD1a+ cells) in whole sections are presented (top). Quantification of double positive cells in each condition was represented as mean ± SD (bottom left), as well as the distribution of double positive cells between the epidermis and the dermis (bottom right). **p < 0.01, as determined by Student’s t-test. ns = non-significant. **(C)** Epidermal cells were purified from skin explants and infected with USUV AF2, WNV L1 at MOI 2 for 24 h. Cells were gated as indicated in Supp. Figure 1C and viral replication was estimated by flow cytometry by staining intracellular viral dsRNA both in CD207+ Langerhans cells (eLCs) and in HLA-DR-/CD207-epidermal cells (mainly keratinocytes). **(D)** Epidermal cells were purified from skin explants and infected with USUV AF2, WNV L1 at MOI 2. At 24 hpi, cells were fixed and stained for nuclei (blue), dsRNA (green) and CD207 (red). Images were acquired on a Leica SP5-SMD microscope. Scale bar: 10 μm. **(E)** Epidermal CD1a+ cells were infected with USUV AF2, WNV L1 at MOI 2 for 24h or 48h. Intracellular viral RNA was quantified by RT-qPCR. Results from 3 independent experiments performed in triplicate are shown. ***p < 0.001, ****p < 0.0001, as determined by Student’s t-test.

### USUV replicates at higher rates than WNV in monocyte-derived LCs

To further investigate the propensity of USUV to infect human LCs, we moved to a model of human monocyte-derived cells. Since monocytes can be differentiated into either DCs (MoDCs) or LCs (MoLCs) (Supp. Figure 1A), we first compared the kinetics of USUV and WNV replication in these cells. In order to exclude any strain-specific phenotype, we included 2 more viral lineages in our study: USUV Europe 2 (EU2, TE20421/Italy/2017), a lineage that has been involved in several severe clinical cases ^40,41^, and a strain of WNV L2 (WNV-6125/France/2018), a lineage that actively circulates in Europe since 2004 ^42^. We infected monocytes, or autologous MoDCs or MoLCs, and infection was followed over time by RT-qPCR. WNV strains showed slow and low increases in the amount of viral RNA over time that were comparable in the 3 cell types, whereas USUV viral loads increased much faster and higher in amplitude in MoLCs and MoDCs, and peaked at 16 and 24 h post-infection (hpi), respectively (Figure 2A). Interestingly, USUV EU2 was the most virulent strain in these 2 models, in terms of kinetic and replication rates.

**Figure 2.**
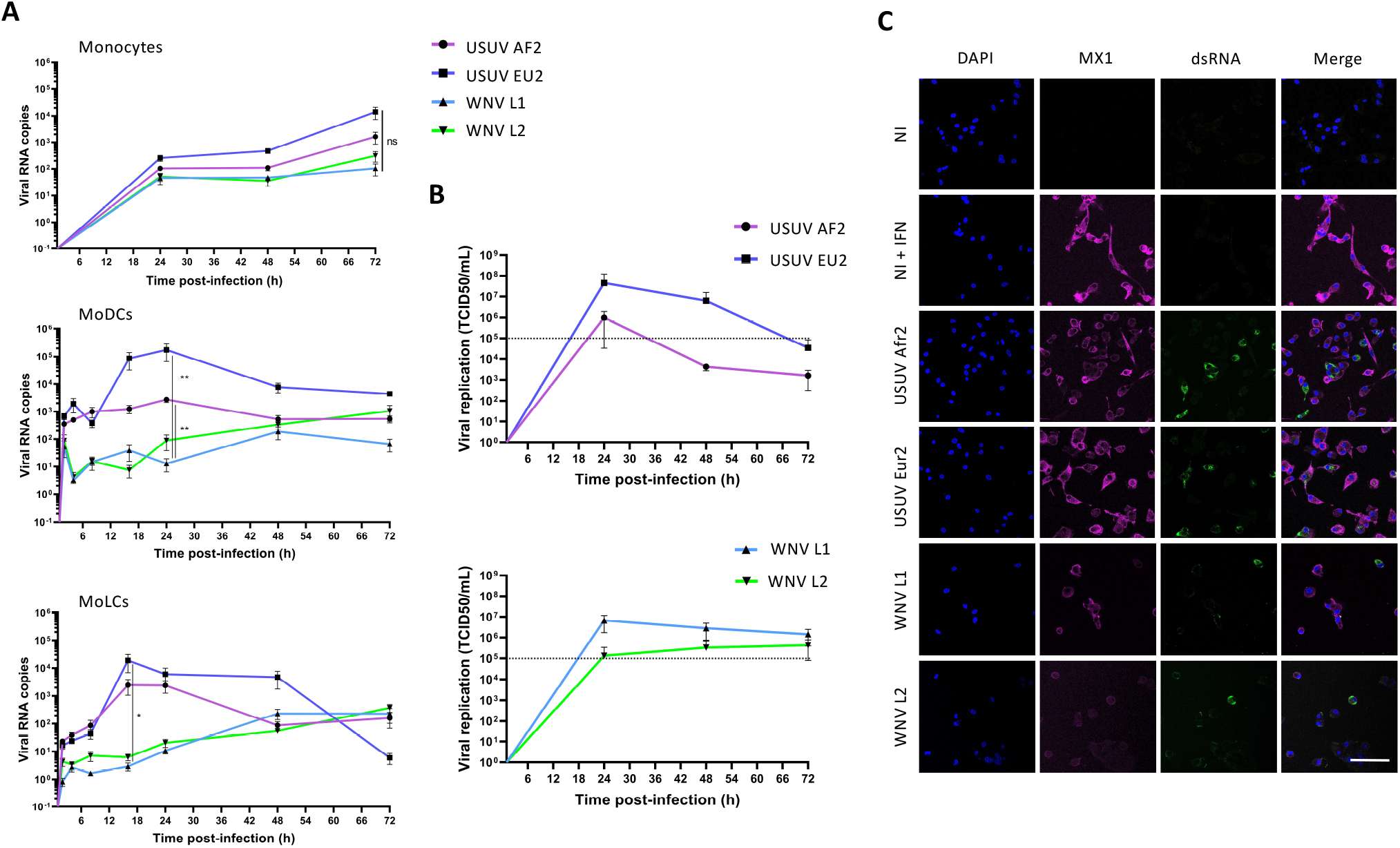
Viral replication of WNV and USUV in primary human myeloid cells. **(A)** Human monocytes and autologous MoDCs or MoLCs were infected by USUV AF2, USUV EU2, WNV L1 or WNV L2 at MOI 1. Cells were harvested at indicated times post-infection and intracellular viral RNA was quantified by RT-qPCR. Data represent 3 independent experiments run in technical duplicate, represented as mean ± SEM and shown in intracellular viral RNA copies. *p < 0.05, **p < 0.01 have been determined using a Mann-Whitney *t*-test relative to indicated condition. **(B)** MoLCs were infected by USUV AF2, USUV EU2, WNV L1 or WNV L2 at MOI 1. Viral production in MoLCs was assessed by TCID50/ml titration of supernatants at indicated times on Vero cells. The dotted line shows the titer of the viral inoculum. Results are represented as mean ± SD of 3 independent experiments performed in duplicate. **(C)** MoLCs were infected at MOI 1 by USUV AF2, USUV EU2, WNV L1, WNV L2, not infected (NI) or pre-treated by IFN-α2 (NI + IFN). Cells were fixed at 24hpi (USUV) or 48 hpi (WNV) and stained for nuclei (blue), dsRNA (green) and MX1 (purple). Images were acquired on a Leica SP5-SMD microscope. Scale bar: 35 µm.

In order to confirm that MoLCs are able to support productive infection of USUV and WNV, we titrated the viruses produced by MoLCs over time. As shown in Figure 2B, replicative virus was detected in MoLC culture medium as early as 24 hpi and the highest viral production rate was again found for USUV EU2, which reached 4.10^7^ TCID50/ml. Noteworthy, the viral replication kinetics of USUV strains showed bell-shaped curves, typical of an effective viral replication (Figure 2B, top), while WNV L1 and WNV L2 RNA copies were significantly much lower and showed no active replication (Figure 2B, bottom). Similarly, immunofluorescence labeling of dsRNA in MoLCs infected with all 4 viral strains in order to evaluate the viral replication showed higher replication levels for USUV strains compared with WNV (Figure 2C). We also looked at Mx1 expression, a type I interferon-induced protein, in order to evaluate the innate immune response of MoLCs to infection ^43^. Interestingly, we observed that dsRNA and Mx1 staining were mutually exclusive, a pattern that was particularly pronounced in the case of USUV (Figure 2C), suggesting that MoLCs respond to USUV infection by secreting interferon, thus partially inhibiting viral replication (Figure 2C). In accord with this, we found that type I IFN strongly inhibits the replication of all 4 viral strains both in MoDCs and in MoLCs (Supp. Figure 2).

### USUV induces a higher innate immune response than WNV in MoLCs

To further investigate the innate immune response of MoLCs to WNV and USUV infection, we first compared the production of type I IFN triggered by WNV and USUV in monocytes, MoDCs and MoLCs. In monocytes, only USUV EU2 infection led to a very limited amount of type I IFN secreted in the culture medium from 48 hpi (Figure 3A). In MoDCs and MoLCs however, high amounts of type I IFN were produced, especially following USUV infection (Figure 3A). The divergence between USUV and WNV infection was particularly striking in MoLCs, since both USUV strains triggered a fast and potent IFN secretion of up to 10^8^ U/ml at 24 hpi (Figure 3A), whereas WNV L1 and WNV L2 induced a delayed response, or no response, respectively. Therefore, we observed a good correlation between the susceptibility to infection of MoLCs by a given virus (Figure 2A) and their propensity to secrete IFN (Figure 3A). Next, we investigated whether USUV triggers a globally more intense antiviral innate immune response than WNV in MoLCs. To this end, we quantified the expression of a large panel of cytokines, chemokines and interferon-stimulated genes (ISGs) by RT-qPCR in MoLCs at 24 hpi. As anticipated, the induction of all transcripts was higher in USUV-compared to WNV-infected cells and once again, USUV EU2 was the most potent trigger (Figure 3B). A selection of cytokines was also quantified at the protein level in the culture medium of MoLCs at 48 hpi, and confirmed the former observations (Figure 3C). Therefore, our results suggest that the efficient replication of USUV in MoLCs triggers a fast and potent innate immune response.

**Figure 3.**
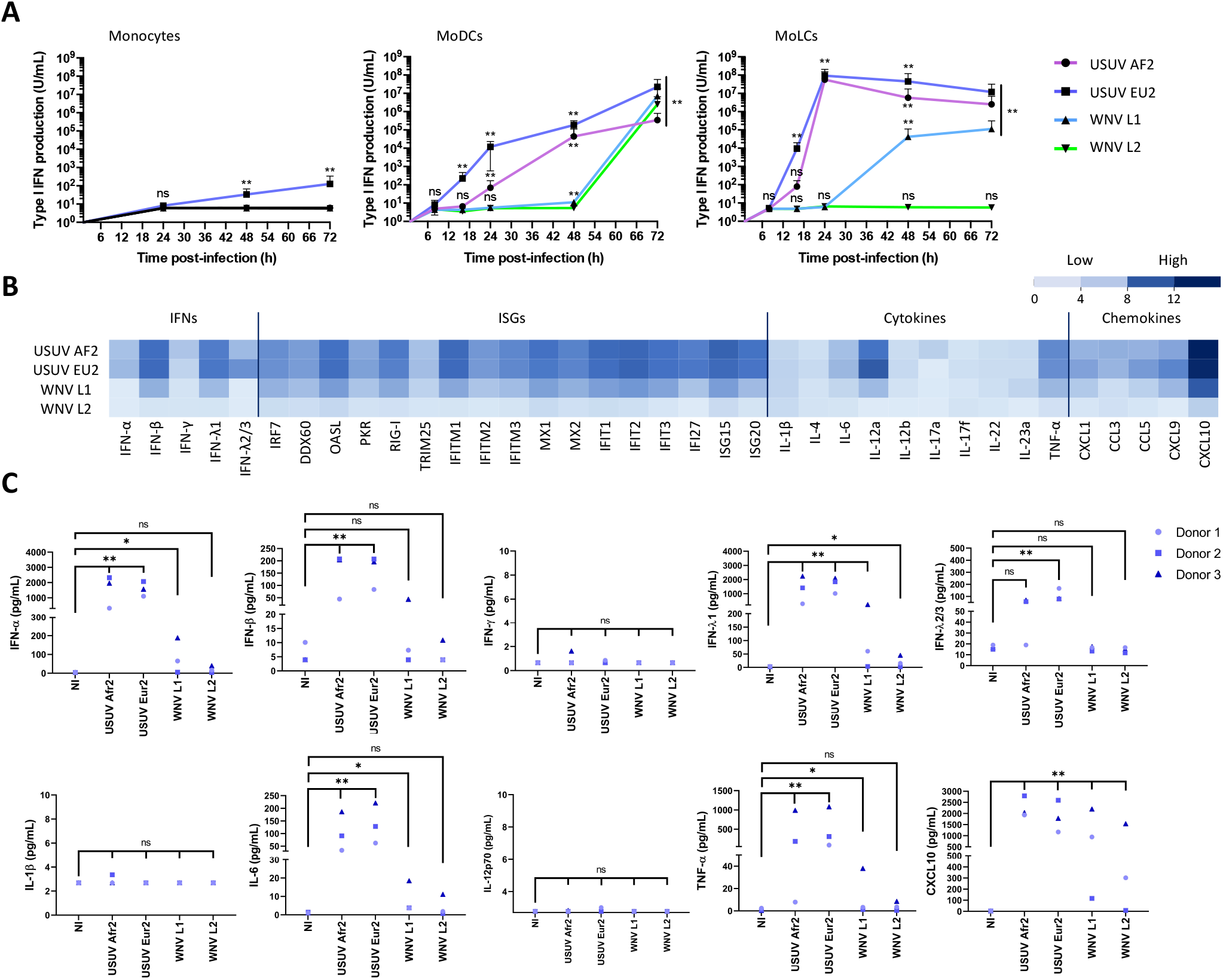
Characterization of USUV- and WNV-triggered innate immune response in MoLCs. **(A)** Human monocytes and autologous MoDCs or MoLCs were infected with USUV AF2, USUV EU2, WNV L1 or WNV L2 at MOI 1 for 8, 16, 24, 48 or 72 hours. Type I IFN secreted in the culture medium by infected cells was quantified on STING-37 reporter cells. Data are from 3 independent experiments performed in duplicate and are represented as mean ± SD. *p < 0.05, **p < 0.01 have been determined using a Mann-Whitney *t*-test relative to control cells. ns = non-significant. **(B)** MoLCs were harvested at 24 hpi and transcripts of a panel of proteins including IFNs, ISGs, cytokines and chemokines were quantified by RT-qPCR. Results are summarized on a heatmap showing low (light blue) to high (dark blue) transcription intensity. Data represent the mean of the log2 transcription intensity of each indicated transcript, from four independent experiments run in technical duplicate, relative to control cells (NI). **(C)** Cytokine concentrations in supernatants from infected MoLCs were measured using the multiplex bead-based immunoassay LEGENDplex (BioLegend) at 48 hpi. Data are from 3 independent experiments performed on cells from 3 donors. *p < 0.05, **p < 0.01 have been determined using a Mann-Whitney *t*-test relative to uninfected cells (NI). ns = non-significant.

### USUV escapes post-entry langerin-mediated restriction

Since we showed that USUV can infect DCs and has a strong tropism for both MoLCs and epidermal LCs, we sought to decipher whether it has a better propensity to use DC-expressed CLR as entry receptors compared with WNV. To do this, we ectopically expressed human DC-SIGN or langerin in HEK293T cells (Supp. Figure 3A), which are poorly permissive to USUV and WNV at low MOI (Figures 4A and 4B), and evaluated both viral entry and replication. At 48 h post-infection, the expression of DC-SIGN enhanced WNV infection by more than 6 times, in agreement with previous studies ^22,32^, whereas the expression of langerin did not allow the entry and/or replication of WNV in HEK293T cells (Figure 4A). In contrast, both DC-SIGN and langerin promoted infection by USUV (Figure 4A), thus suggesting that unlike WNV, USUV can also use langerin as a receptor to enter and replicate. These observations were confirmed by quantifying the amount of viral RNA in infected cells by RT-qPCR, since we observed that DC-SIGN expression led to a 10-fold increase of both USUV and WNV infection, whereas langerin expression only increased USUV replication (Figure 4B). We also confirmed by western-blot that langerin expression allowed a marked enhancement of USUV E protein detection in infected cells at 24 h and 48 h (Figure 4C and Supp. Figure 3B). In the case of WNV, again, langerin expression had either no effect on virus replication or was even deleterious (Figure 4C). These results pointed to two possible interpretations: on the one hand, this could mean that only USUV can use langerin as a receptor, on the other hand, it is possible that both viruses enter via langerin, but only USUV manages to escape langerin-induced degradation and replicate. To distinguish between these two hypotheses, we tested the ability of langerin to interact with USUV and WNV envelope proteins by co-immunoprecipitation. To this end, we incubated langerin-overexpressing HEK293T cells (Supp. Figure 3C) with USUV or WNV at MOI 5 for 30 minutes. Since langerin recognizes mannose-rich glycans present on viral glycoproteins, we performed competition experiments using mannan [15,24]. Our results indicate that langerin can interact with both USUV and WNV envelope proteins and that, as expected, this interaction can be inhibited by mannan (Figure 4D). Consistent with the role of langerin in pathogen recognition and capture, we observed by FACS and microscopy that langerin not only binds USUV and WNV, but also allows their internalization (Figures 4E and 4F). Taken together, our results suggest that langerin is able to recognize and internalize both USUV and WNV, but that only USUV is able to subsequently replicate.

**Figure 4.**
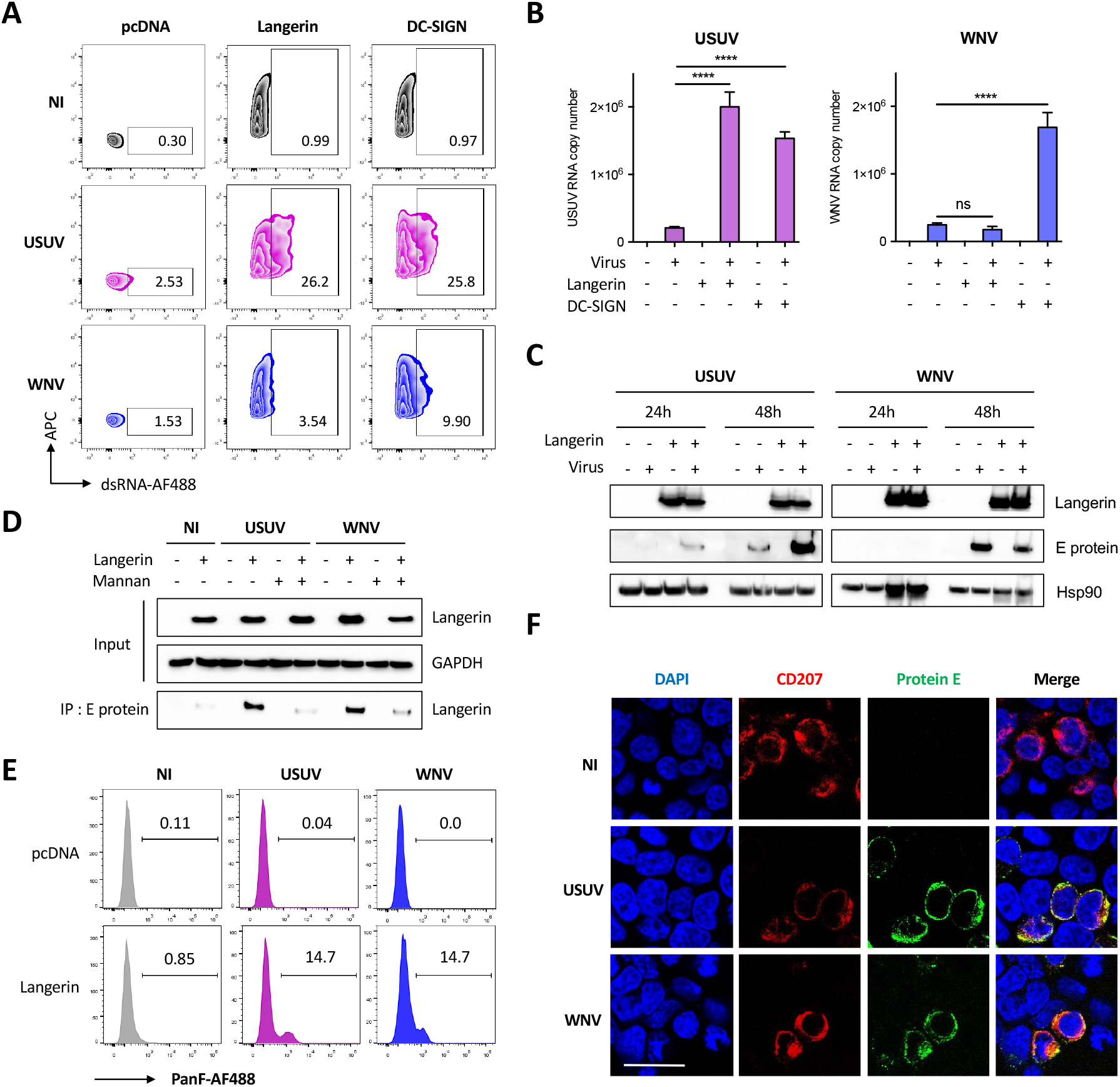
Langerin expression allows USUV but not WNV replication in HEK293T cells. (**A** and **B**) HEK293T cells were transfected with an empty plasmid (pcDNA) or with plasmids encoding langerin or DC-SIGN. 24h post transfection, cells were infected with USUV AF2 and WNV L1 at MOI 0.5 for 48 h, or left non-infected (NI). (**A**) USUV and WNV replication was assessed by flow cytometry following intracellular immunostaining using anti-dsRNA and Alexa Fluor 488, anti-langerin-APC and anti-DC-SIGN-APC antibodies. The percentage of infected cells among langerin-positive and DC-SIGN-positive (transfected) cells is shown. Results are from one representative experiment of 3 independent experiment. **(B)** Same as (**A**), except that viral replication was estimated by quantifying viral RNA by RT-qPCR. Results are represented as mean fold change ± SD of 3 independent experiments performed in duplicate, and are expressed as viral RNA copies. ****p < 0.0001, as determined by Student’s t-test. ns = non-significant. **(C)** HEK293T cells were transfected with an empty plasmid (-) or with a plasmid encoding human langerin (+), as indicated. At 24 h post transfection, cells were infected with USUV AF2 or WNV L1 at MOI 0.5 for 24 h or 48 h, or left uninfected. Cell lysates were subjected to western-blotting using anti-langerin or pan-flavivirus antibodies. Quantification of Western Blot analyses is shown in Supp. Figure 3B. **(D)** HEK293T cells were transfected or not with a plasmid encoding langerin. At 24 h post transfection, cells were pretreated or not with 1 mg/ml mannan for 1 h and infected with USUV AF2 and WNV L1 at MOI 5 for 30 min, or left noninfected (NI). Following cell lysis, whole-cell lysates were subjected to IP with anti-E protein antibody followed by Western blot analysis with the anti-langerin antibody. **(E)** HEK293T cells were transfected with an empty plasmid (pcDNA) or with a plasmid encoding langerin. At 24 h post transfection, cells were infected with USUV AF2 and WNV L1 at MOI 5 for 30 min, or left noninfected (NI). Cells were fixed and subjected to flow cytometry following surface immunostaining the viral E protein with the pan-flavivirus anti-E antibody. The percentage of cells expressing the viral E protein is shown. Results are from one representative experiment of 2 independent experiments. **(F)** HEK293T cells expressing langerin were left uninfected (NI) or infected by USUV AF2 or WNV L1 at MOI 5 for 30 min. Cells were fixed and stained for nuclei (blue), langerin (red) and E protein (green). Images were acquired on a Leica SP5-SMD microscope. Scale bar: 15 µm.

In order to confirm our observations in primary cells, we performed further experiments using MoLCs and epidermal LCs. We confirmed that mannan efficiently prevented the infection of LCs by USUV, as shown by RT-qPCR amplification of the viral genome (Figure 5A) and flow cytometry using the dsRNA antibody (Figure 5B). Moreover, we silenced langerin expression in eLCs purified form human epidermis using a specific siRNA (Figure 5C) and show that this efficiently reduced infection by USUV (Figure 5D), thus formally demonstrating that langerin is an entry receptor for USUV in human LCs. As observed in HEK293T cells overexpressing langerin (Figure 4F), we noted that endogenous langerin co-localized with incoming USUV in the very first steps of infection, thus suggesting that langerin is co-internalized with USUV virions in eLCs (Figure 5E).

**Figure 5.**
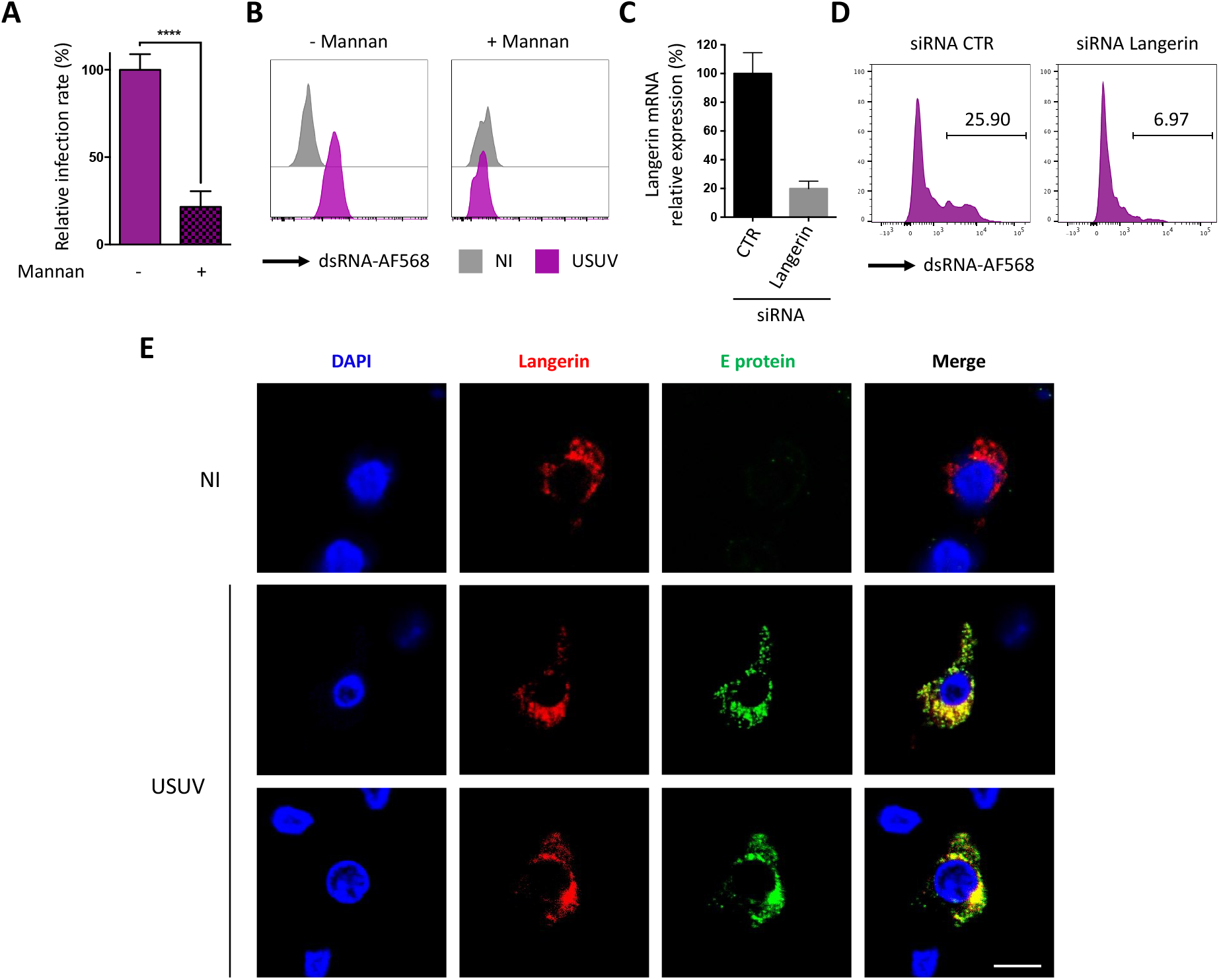
Langerin allows USUV entry and replication in human LCs. **(A)** MoLCs were left untreated or treated for 1 h with 1 mg/ml mannan and infected with USUV AF2 at MOI 2 for 24 h. Viral replication was assessed by quantifying the amount of viral RNA in cells by RT-qPCR. Data represent the mean ± SD of 3 independent experiments performed in triplicate on cells from 3 different donors. **(B)** Epidermal LCs purified from human skin were treated or not with 1 mg/ml mannan for 1 h and infected with USUV AF2 at MOI 2. At 24 hpi, viral replication was estimated by flow cytometry using dsRNA and Alexa Fluor 568 antibodies. Data are from one representative experiment of two independent experiments. (**C** and **D**) Epidermal LCs purified from human skin were transfected with siRNA control (CTR) or siRNA targeting langerin. Relative expression of langerin was estimated by RT-qPCR analysis (**C**). Cells were infected with USUV AF2 for 24 h at MOI 2 and viral replication was assessed by flow cytometry using anti-dsRNA and AF568 antibodies (**D**). Data are from one representative experiment of two independent experiments. (**E**) Epidermal LCs purified from skin explants were infected (USUV) or not (NI) with USUV AF2 for 30 min at MOI 5. Cells were fixed and stained for nuclei (blue), E protein (green) and CD207 (red). Images were acquired on a Leica SP5-SMD microscope. Scale bar: 20 µm.

## Discussion

Arboviruses such as Dengue, Zika, West Nile, or Usutu viruses represent a global public health threat due to globalization and worldwide spread of mosquito vectors ^5,44^. Since mosquito-borne viruses are directly inoculated in the epidermis and the dermis during blood meals, the skin constitutes the initial site of viral replication and immune response ^28^. LCs and dDCs patrol the epidermis and the dermis, respectively, to sense and capture pathogens. For this purpose, they are equipped with unique receptors, known as CLRs, which bind carbohydrate moieties associated to pathogens. Among them, DC-SIGN and langerin are expressed by DCs and LCs, respectively, and act both as PRRs and antigen-uptake receptors. Therefore, DC-SIGN and langerin constitute key receptors, allowing the interception of a variety of pathogens that enter the organism through the skin ^12,45^. In the case of viruses, glycans are present on their envelope glycoproteins, which are recognized by CLRs, thus inducing their endocytosis and subsequent degradation. But like many cellular defense mechanisms, DC-SIGN and langerin can be bypassed or even hijacked by some viruses to their advantage. Thus, many viruses can use DC-SIGN as a receptor to propagate, either in *cis*, via the productive infection of DCs, or in *trans*, if the receptor facilitates the capture and transmission to other cells ^12,18^. In agreement with this, we found that both USUV and WNV can use DC-SIGN as a receptor, as already shown for many flaviviruses ^21,32^.

In contrast to DC-SIGN, hijacking of langerin by viruses seems much rarer, and has so far only been demonstrated for IAV in transfected cell lines ^24^. In this manuscript, we show that USUV, but not WNV, can use naturally expressed langerin to infect into LCs. Our results not only show that LCs are permissive to USUV, but also that they support productive viral replication. This is a surprising observation, since LCs are notoriously refractory to most viruses, with only a few exceptions, including HSV-1 ^46,47^ and DENV ^25–27^. Among the viral strains that we tested, USUV EU2 showed the fastest and most efficient replication in both DCs and LCs, and also triggered the most intense innate immune response. Interestingly, this viral strain was previously described as being involved in several clinical cases, and was recently shown to be particularly neurovirulent and lethal in mice ^40,41,48^

Flaviviruses encode one, two, or no N-linked glycosylation sites on their envelope proteins (E protein) ^23^. In the case of WNV, it was shown that, unlike non-glycosylated viral particles, glycosylated strains can use DC-SIGN to infect DCs ^32^, thus illustrating the importance of N-glycosylations for flavivirus tropism. WNV and USUV E proteins contain a single N-linked glycosylation site at residue 154, whereas most DENV isolates contain glycosylation sites at residues 153 and 67 ^49,50^. The specificity of the interactions between glycoproteins and CLRs is complex and depends both on the type and position of glycans. For example, whereas WNV grown in mammalian cells was shown to preferentially use DC-SIGNL as a receptor ^22^, the introduction of a glycosylation site at position 67 into WNV E protein conferred the capacity to also use DC-SIGN ^51^. From the sequence of its USUV E protein, it can be predicted that USUV has, like WNV, a unique glycosylation at N154 ^48^, thus explaining why both viruses can interact with langerin.

In the case of USUV, we showed that langerin expression not only allows the virus to bind and enter, but also to replicate, thus suggesting that in this case, langerin could be considered as a receptor. However, further work will be required in order to determine whether langerin acts as a *bona fide* entry receptor or as a proviral factor facilitating virion attachment and entry. In the case of DENV for instance, it was shown that endocytosis-defective DC-SIGN allows viral entry as efficiently as the wild type protein, thus suggesting that DC-SIGN is likely an attachment factor rather than an entry receptor ^52^. In our case however, we showed that blocking or silencing langerin in LCs prevents USUV infection, whereas the ectopic expression of langerin in non-permissive cells promotes virus binding, entry and replication. Thus, it is likely that langerin is necessary and sufficient to allow USUV infection of LCs. Furthermore, we show that, although WNV also binds langerin and is internalized in langerin-expressing cells, this does not allow its replication, presumably because the virus is degraded in Birbeck granules, as demonstrated in the case of HIV-1 ^15^. Since USUV and WNV are phylogenetically closely related ^1,2^, our results suggest that adaptation of USUV to human cells involved bypassing langerin-mediated degradation to infect skin-resident Langerhans cells. Further work will be required to uncover the replication advantage that this brings to USUV, in particular whether it contributes more efficiently to its dissemination.

As the most peripheral immune cells in the body, LCs are the most exposed. In this respect, it is rather surprising that so few viruses have been found to infect them. It is possible that langerin is more difficult than DC-SIGN to be hijacked by viruses, especially since it forms Birbeck granules that efficiently degrade viruses ^13–15^. Alternatively, this may just reflect the fact that LCs are less studied than other types of dendritic cells. In any case, langerin has been described as a potent antiviral barrier that very few viruses are able to overcome ^15,24,25,47,53^. How USUV, once inside LCs, manages to avoid degradation in Birbeck granules to replicate in LCs remains an open question.

Finally, our results showed a correlation between the susceptibility of LCs to infection and their ability to respond to this infection. Indeed, USUV strains, which infect LCs more efficiently than WNV, also induce a stronger and faster innate immune response. Similarly, among the two USUV strains tested, EU2, the better and fastest replicating strain, was also a more potent inducer of antiviral cytokines and chemokines, including type I IFN. It is intriguing that this virus can replicate so efficiently while inducing such an intense innate response. Viruses are in a speed race with the IFN response to replicate before the IFN-induced antiviral state is established and our results suggest that USUV EU2 is fit enough to win this race. USUV was previously described to induce a stronger interferon response than WNV in MoDCs ^54^. Interestingly, authors also showed that USUV was more sensitive than WNV to the antiviral activity of types I and III IFNs ^54^. This latter observation might explain why USUV replicates faster than WNV, in order to overtake cellular defenses by completing their replication cycle before the expression of antiviral ISGs. ISGs interfering with WNV replication are now well characterized ^55^, but no study has yet been performed on USUV, thus illustrating the fact that research on USUV is still in its infancy. A number of studies, including our own, highlight important differences between USUV and WNV in terms of virus-cell and virus-host interactions ^54,56^. Since it is therefore impossible to transpose our knowledge of WNV onto USUV, more research is clearly needed in order to anticipate the possible worldwide emergence of this virus and the burden to economy and public health it may pose in the future.

## Acknowledgments

We acknowledge the imaging facility MRI (Montpellier, France), member of the national infrastructure France-BioImaging, supported by the French National Research Agency (ANR-10-INBS-04, “Investissements d’avenir” program). We also thank Dr Alain Fautrel from the H2P2 platform (Université de Rennes 1, Rennes, France) for his help and his reactivity.

This work was supported by the Labex EpiGenMed, an “Investissements d’Avenir” program (ANR-10-LABX-12-01). M.F.M was the recipient of a doctoral fellowship from the Labex EpiGenMed. G.M. was supported by a grant from the Agence National de la Recherche sur le SIDA et les Hépatites virales (ANRS).

## Supp Tables

**Supp Table 1.**
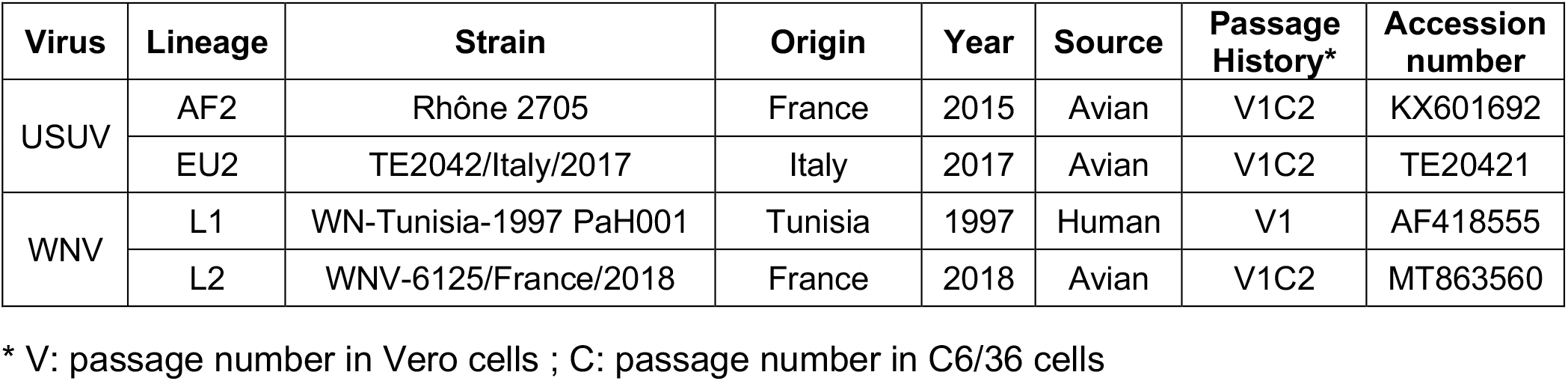
Origin and history of isolates used in the study.

| Virus | Lineage | Strain | Origin | Year | Source | Passage History* | Accession number |
| --- | --- | --- | --- | --- | --- | --- | --- |
| USUV | AF2 | Rhône 2705 | France | 2015 | Avian | V1C2 | KX601692 |
|  | EU2 | TE2042/Italy/2017 | Italy | 2017 | Avian | V1C2 | TE20421 |
| WNV | L1 | WN-Tunisia-1997 PaH001 | Tunisia | 1997 | Human | V1 | AF418555 |
|  | L2 | WNV-6125/France/2018 | France | 2018 | Avian | V1C2 | MT863560 |

\* V: passage number in Vero cells ; C: passage number in C6/36 cells

**Supp Table 2.**
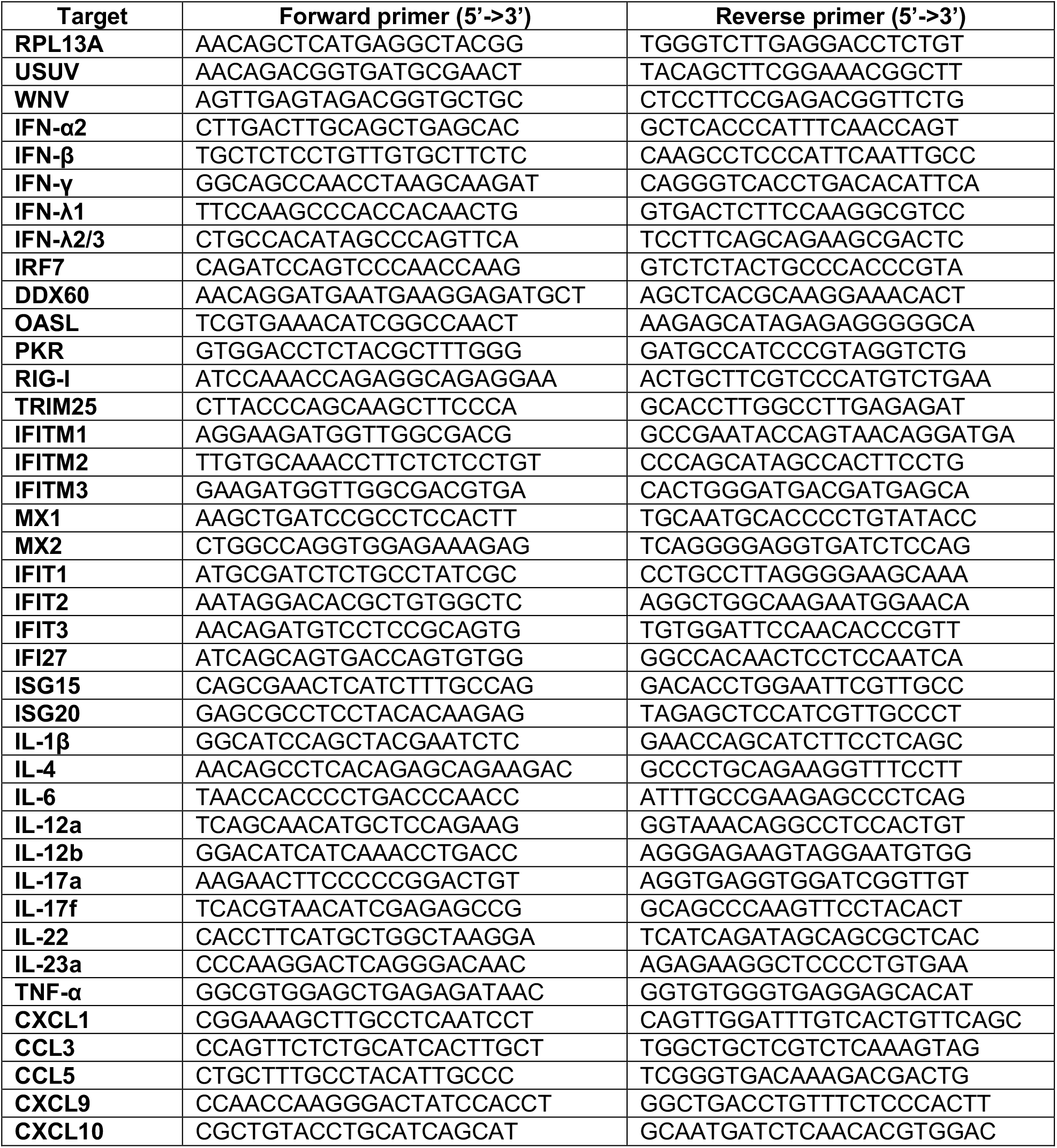
Primers used in RT-qPCR analyses.

## Supp Figures

**Supp. Figure 1.**
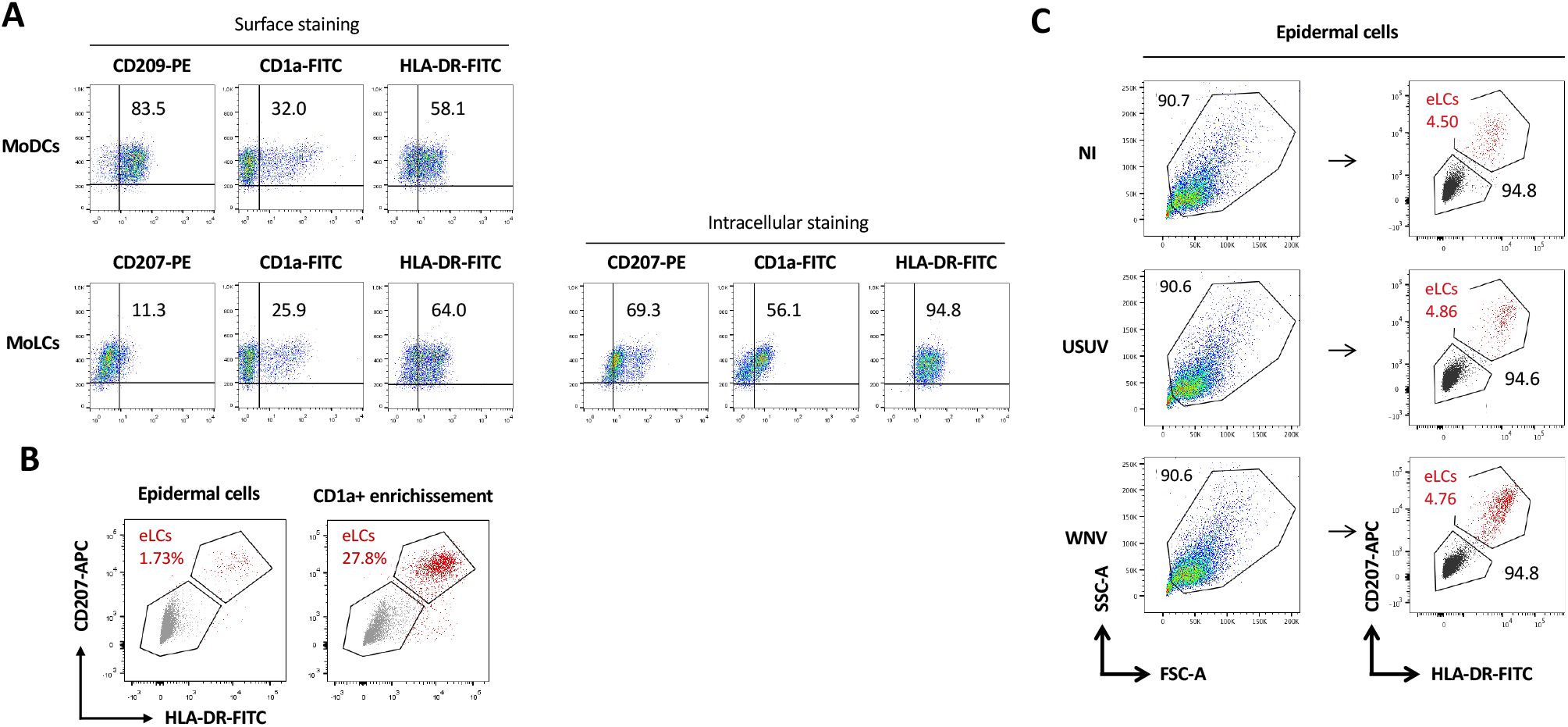
Phenotyping of epidermal cells, eLCs, MoDCs and MoLCs. (**A**) Expression of CD209 (DC-SIGN), CD207 (Langerin), CD1a and HLA-DR (MHC class II) was assessed by flow cytometry using appropriate antibodies. MoDCs were subjected to surface staining only (top panel) and MoLCs to both surface (bottom left panel) and intracellular (bottom right panel) staining. Positive cells for each marker appear in the top right quarter of each dot-plot. Representative phenotypes are shown. **(B** and **C)** Expression of CD207 and HLA-DR (MHC class II) was assessed by flow cytometry using anti-CD207-APC and anti-HLA-DR-FITC (Class-II-FITC) antibodies, respectively. HLA-DR+/CD207+ Langerhans cells (eLCs) are colored in red and HLA-DR-/CD207-epidermal cells (mainly keratinocytes) appear in grey. A representative phenotyping is shown for B and C. **(B)** Staining was performed both on total epidermal cells (left panel) and following enrichment of CD1a+ cells (right panel). **(C)** Gating strategy for analyzing non-infected and infected epidermal cells, related to Figure 1C. Cells were gated in an FSC-A and SSC-A dot plot, before gating eLCs (HLA-DR+/CD207+) and HLA-DR-/CD207-epidermal cells. Both populations were analyzed in Figure 1C.

**Supp. Figure 2.**
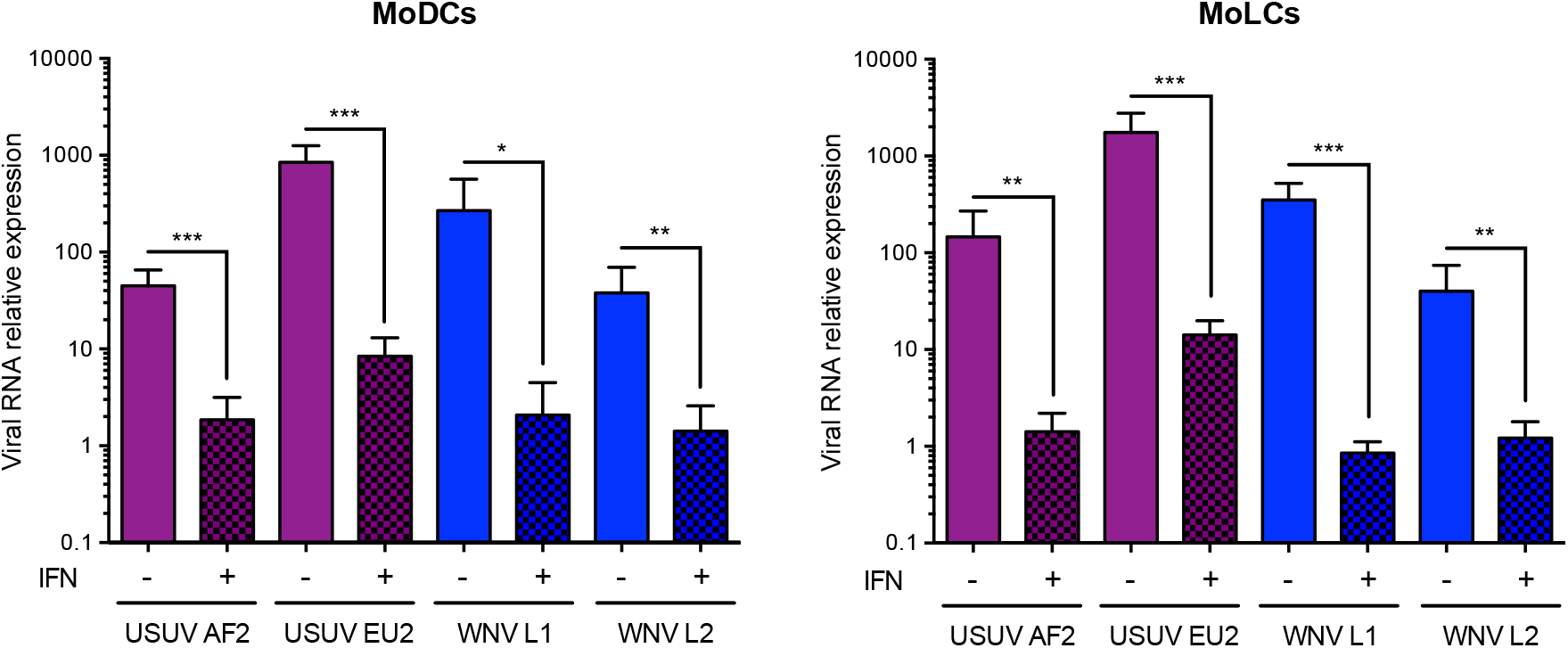
IFN-α2 inhibits USUV and WNV replication in MoDCs and MoLCs. MoDCs or MoLCs were pre-treated or not with 1000 U/mL of IFN-alpha2 for 24 h before infection with USUV AF2, USUV EU2, WNV L1 or WNV L2 at MOI 1. At 24 hpi, total RNA were extracted and viral RNA was quantified by RT-qPCR. Results from 3 independent experiments performed in duplicate are shown. ***p < 0.001, **p < 0.01, *p < 0.05, as determined by Student’s t-test.

**Supp Figure 3.**
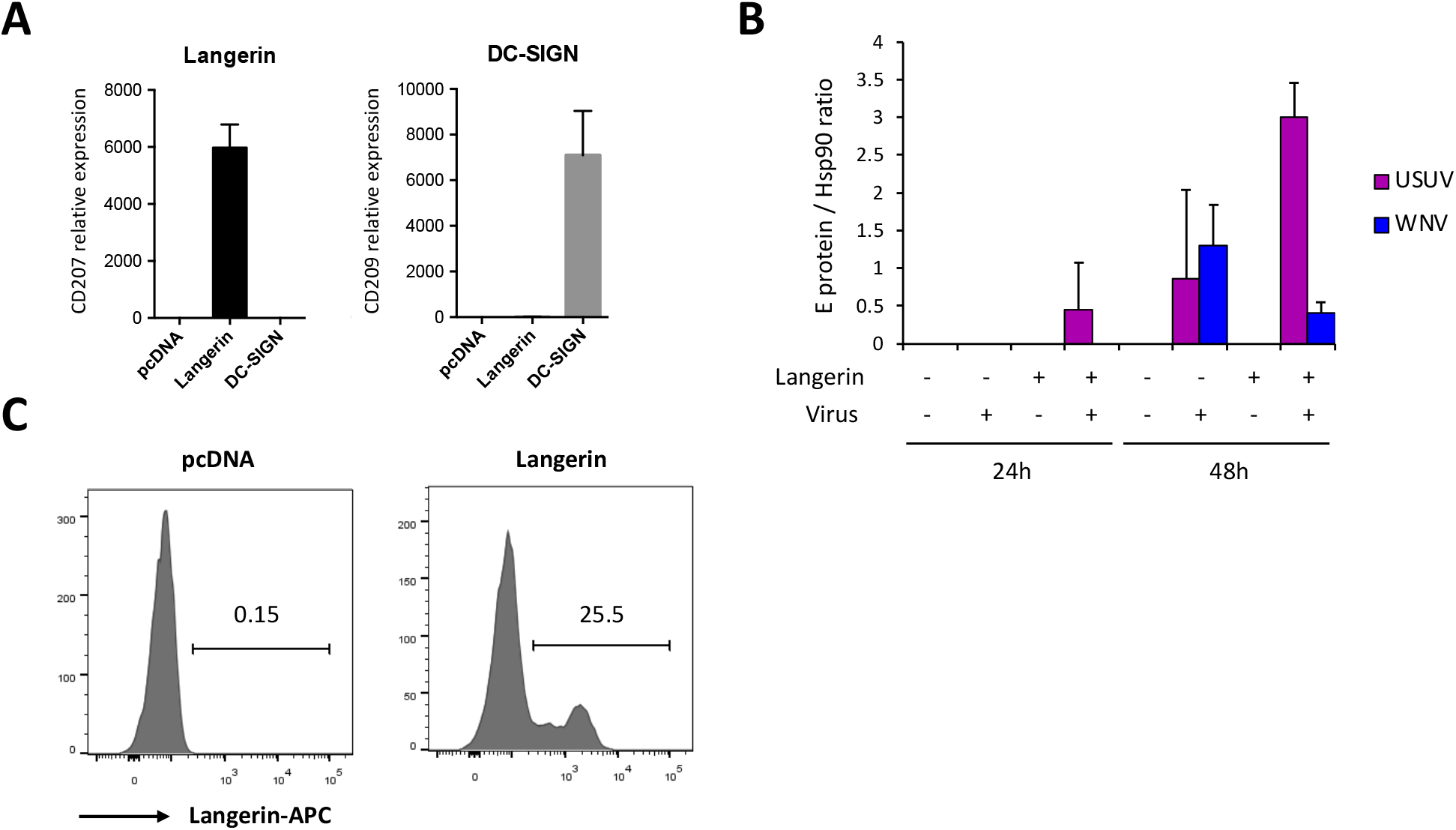
Expression levels of Langerin, DC-SIGN and viral E protein in HEK293T cells. **(A)** In parallel of the experiments shown in Figures 4A and 4B, the expression of Langerin and DC-SIGN in HEK293T cells was estimated at 24 h post-transfection by RT-qPCR. Relative expression of Langerin and DC-SIGN mRNA expression is represented as mean values ± SD. **(B)** Quantification of western-blots presented in Figure 4C. The intensity of the E protein and Hsp90 bands was quantified using the Image Lab Software (Bio-Rad Laboratories). The mean ± SD of the E protein/Hsp90 ratio is represented (n=2). **(C)** The expression of Langerin in transfected HEK293T cells used in the experiment shown in Figure 4D and 4E was evaluated by flow cytometry using an anti-langerin antibody. The percentage of Langerin-expressing cells is indicated.

